# Comparison of Y-chromosomal lineage dating using either evolutionary or genealogical Y-STR mutation rates

**DOI:** 10.1101/004705

**Authors:** Chuan-Chao Wang, Hui Li

## Abstract

We have compared the Y chromosomal lineage dating between sequence data and commonly used Y-SNP plus Y-STR data. The coalescent times estimated using evolutionary Y-STR mutation rates correspond best with sequence-based dating when the lineages include the most ancient haplogroup A individuals. However, the times using slow mutated STR markers with genealogical rates fit well with sequence-based estimates in main lineages, such as haplogroup CT, DE, K, NO, IJ, P, E, C, I, J, N, O, and R. In addition, genealogical rates lead to more plausible time estimates for Neolithic coalescent sublineages compared with sequence-based dating.

## Introduction

The paternally inherited Y chromosome has been widely used in anthropology and population genetics to understand demographic history of human populations (Wang and Li, 2013). There are two kinds of extremely useful markers in Y chromosome, single nucleotide polymorphism (SNP) and short tandem repeat (STR). Over the last two decades, SNP and STR have been widely used in Y-chromosomal diversity studies (Jobling and Tyler-Smith, 2003). The most important link between genetic diversity and human history is time, for instance, the time when a lineage originated or expanded, or when a population split from another and migrated. Y-STR has also been used in time estimation for SNP lineages. Although this approach is widely used, there are still many ongoing debates about the best way to use STRs in lineage dating. In particular, there are two popularly used Y chromosomal STR mutation rates, that is, the genealogical rate and the evolutionary rate. The genealogical rates are directly observed rates in deep-rooted pedigrees or father-son pairs (Wei et al., 2013a; Zhivotovsky et al., 2004). The evolutionary rates are those calibrated against historical events, such as the divergence of the Maoris and Cook Islanders in the Pacific (Zhivotovsky et al., 2004). To choose which kind of mutation rate in the Y chromosome dating is controversial, since different rates can result in several-fold deviation.

With the advent of next-generation sequencing technology, Y chromosomes of numerous human individuals have been entirely sequenced recently (Wei et al., 2013b; Poznik et al., 2013; Francalacci et al., 2013; Yan et al., 2013; 1000 Genomes Project Consortium, 2012). The increasing large amount of Y chromosomal sequence data provides a very good opportunity to evaluate the previously proposed different kind of Y-STR mutation rates in time estimation (Wei et al., 2013a). Here, we have compared the Y chromosomal lineage dating between sequence data and commonly used Y-SNP plus Y-STR data using Batwing.

## Materials and methods

The 1000 genomes dataset: About 8.9Mb sequence data on the unique regions of Y chromosome of the 377 male individuals were extracted from the 1000 Genomes Project Phase I from publicly accessible FTP sites (1000 Genomes Project Consortium, 2012) (supplementary table.1). Y chromosomal haplogroups classification (Van Geystelen et al., 2013), maximum likelihood tree construction (Guindon et al., 2010), and divergence time calculation (Drummond et al., 2002; Drummond and Rambaut, 2007) were following our previous works (Yan et al., 2013; Wang et al., 2013a). The STR data is also downloaded from FTP sites of 1000 Genome Project. The 23 STRs are the same as reported in Wei et al (2013). Median-joining networks (Bandelt et al., 1999) of haplotypes consisting of 21 YSTRs and 35 Y-SNPs were constructed using Network 4.6.1.2 (Fluxus Engineering). Li Jin lab dataset: We selected 78 samples from our previous next-generation sequencing dataset, covering most sublineages of Haplogroup O, as well as Haplogroup C, D, G, J, N, Q, and R (Yan et al., 2013). Seventeen Y chromosome STRs were amplified using the AmpFlSTR Yfiler PCR Amplification kit and analyzed (Yan S and Wang CC et al., unpublished data). The details about next generation data analysis, phylogenetic tree calculation, and time estimation have been reported in our previous work (Yan et al., 2013). In addition, 366 male individuals belonging to haplogroup Oγ-F11 from widely distributed East Asian populations were also included in the Batwing analysis (Wang et al., 2013b).

Time estimation for each Y chromosomal lineage were made using BATWING (Wilson et al., 2003) based on Y-SNP plus Y-STR method, under a model of exponential growth from an initially constant-sized population. The parameters used in estimation were following Xue et al (2006). Five sets of Y-STR mutation rates were applied in time estimations as Wei et al did (Wei et al., 2013a). These are a widely used evolutionary mutation rate (EMR) (Zhivotovsky et al., 2004), a recalibrated evolutionary mutation rate (rEMR) (Shi et al., 2010), two observed genealogical mutation rates (OMRB and OMRS) (Burgarella et al., 2011; Shi et al., 2010), and a genealogical mutation rate adjusted for population variation using logistic model (lmMR) (Burgarella et al., 2011). A total of 10^4^ samples of the program’s output representing 10^6^ MCMC cycles were taken after discarding the first 3×10^3^ samples as burn-in. The Time to the Most Recent Common Ancestor (TMRCA) is calculated using the product of the estimated population size N and the height of the tree T (in coalescent units) (Wilson et al., 2003). A generation time of 25 years was used to produce a time estimate in years. Pearson’s correlation coefficient (r), Spearman’s rank correlation coefficient (rho), and their significance were calculated using R-3.0.2 (http://www.r-project.org).

## Results

The 377 male individuals extracted from the 1000 Genomes Project contain haplogroup A, B, C, D, E, G, I, J, N, O, Q, R, and T, and thus give a good representation of worldwide paternal lineages. The topology of maximum likelihood tree of those samples is congruent with the existing human Y chromosome tree (fig.1a, supplementary fig.1). The length of the branch in the tree is proportional to the number of mutations, and therefore also informative about the times when lineages diverged. The branch length between haplogroup A and out-of-Africa primary haplogroup CT is extremely long, implying they have diverged since a very long time ago. A great number of branches have emerged in the near terminal of the tree, which gives a signal of recent population expansion. The within lineage population expansions are also observed in the Y-STR network, especially in lineage R1b1a2a1a2, E1b1a1a1f1a, E1b1a1a1g, O2b, N1c1a1a2a, and I1a1b (fig.1b). However, the Y-STR network fails to reveal the ancient phylogenetic structure correctly. Haplogroup A individual has not been placed in a very long branch in the network as suggested in the maximum likelihood SNP tree. Haplogroup G is grouped with haplogroup C, and haplogroup T is placed in the same branch with Q and R in the network. Haplogroup R branches from haplogroup Q, with the SNP M242 that defines Q being assigned as recurrent. The similar situation has also been observed in haplogroup D and E, I and J in the network tree. As the mutation rates of STR markers are about four to five orders of magnitude higher than SNPs, the sequence-based phylogenetic tree is much more reliable. The obvious inconsistency between sequence-based and STR-based tree remind us that there might be some bias in Y-chromosomal lineage dating using STR data.

**Fig. 1.**
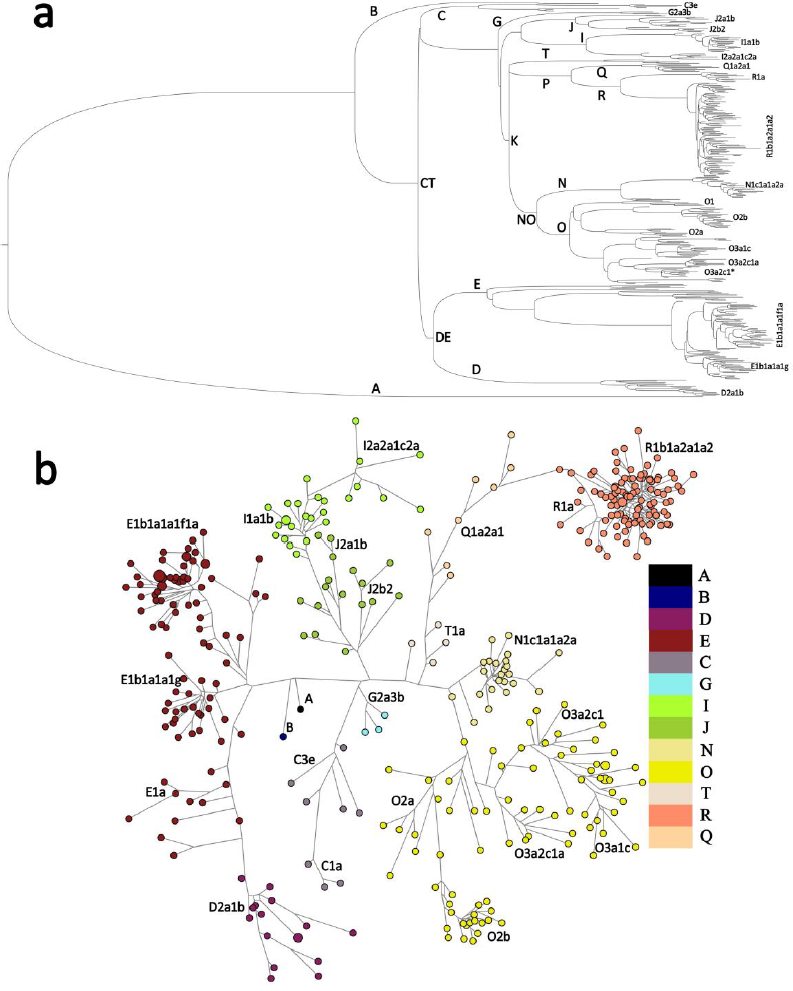
**a.** Phylogenetic tree of human Y chromosome. This tree was constructed using 377 samples sequenced in 1000 Genomes Project. The branch lengths are proportional to the number of SNPs on the branch. For more details, see supplementary fig.1; **b.** Median-joining network representing the relationships between 377 Y chromosomes based on 35 variable Y-SNPs (classified the following haplogroups: A, B, CT, CF, DE, C, C1, C3, D, E, E1a, E1b1a1a1g, E1b1a1a1f, E2, F, G, IJ, I, I1, I2, J, K, NO, N, O, O1, O2, O2b, O3, T, P, R, R1a, R1b, Q) and 21 Y-STRs. Each circle represents a haplotype and has an area proportional to its frequency.

To infer the time depth of Y-chromosomal lineages, we calculated the date of each divergence event throughout the sequence-based tree using Bayesian method. The time to the most recent common ancestor (TMRCA) for all the 377 Y chromosomes estimated was 104.97 thousand years ago (kya) (95% CI: 100.25-109.64 kya). This is consistent with the published estimate of 105 kya (Cruciani et al., 2011) and 101-115 kya (Wei et al., 2013b) for haplogroup A1b1b2b-M219. The next most important split point is the out-of-Africa superhaplogroup CT, which we date here at 56.26 kya (95% CI: 54.29-58.39 kya). This corresponds well to our previous estimation of CT using 78 East Asian Y chromosomes at 3.9 Mbp of the NRY (54.1 kya with 95% CI: 50.6-58.2 kya) (Yan et al., 2013). Only 2 ky later, DE branched off from CT. Most of other main branches (K, NO, IJ, P, E, C, I, J, O, and R) emerged between 20-44 kya. A great number of sublineages branched off from the above main haplogroups in Neolithic time.

We then compared sequence-based time estimation with Y-SNP plus Y-STR based dating. We first used 21 STR markers in Batwing estimation. The TMRCA of all the 377 Y chromosomes estimated using evolutionary STR mutation rates is 117-127 kya, slightly higher than sequence-based TMRCA. However, the estimations using three genealogical mutation rates give the date almost 4-5 times lower than sequence-based TMRCA. This point is consistent with Wei et al’s observation (Wei et al., 2013a). However, the ages for other main lineages (CT, DE, K, NO, IJ, P, E, C, I, J, N, O, and R) show large gaps with both the times estimated using evolutionary and genealogical STR mutation rates. The times using evolutionary rates show a slightly better correlation with the sequence-based estimation than using genealogical rates at the Y chromosomal main lineage level (EMR: Pearson’s r=0.892, Spearman’s rho=0.940, p=1.878E-6; rEMR: Pearson’s r=0.872, Spearman’s rho=0.907, p=1.930E-5; OMRB: Pearson’s r= 0.878, Spearman’s rho=0.923, p=6.852E-6; OMRS: Pearson’s r=0.865, Spearman’s rho=0.896, p=3.481E-5; lmMR: Pearson’s r=0.860, Spearman’s rho=0.879, p=7.545E-5). For the sublineages coalesced in Neolithic Time (C3e, and from D2a1b to R1b1a2a1a2 in x-axis of fig.2a), the TMRCAs based on three genealogical rates are much more consistent with sequence-based TMRCAs than those based on evolutionary rates. At the sublineages level, the ages estimated using genealogical rates have a slightly better correlation with sequence-based estimation (EMR: Pearson’s r=0.651, Spearman’s rho=0.558, p=0.016; rEMR: Pearson’s r=0.652, Spearman’s rho=0.622, p=0.006; OMRB: Pearson’s r=0.688, Spearman’s rho=0.659, p=0.003; OMRS: Pearson’s r=0.715, Spearman’s rho=0.661, p=0.003; lmMR: Pearson’s r=0.649, Spearman’s rho=0.548, p=0.004).

**Fig. 2.**
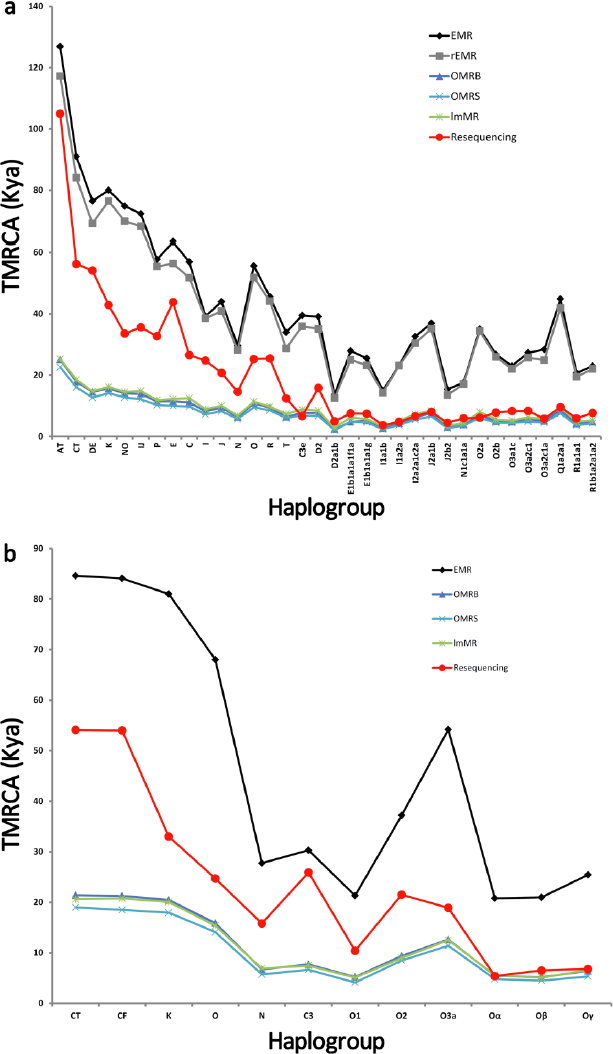
**a.** Comparison of TMRCAs based on Y-SNP and 21 Y-STRs using five different Y-STR mutation rates in 377 samples of 1000 genome project, with the dates estimated based on sequence data. The duplicated locus DYS385 was not used in these analyses, and DYS389 was treated as DYS389I and DYS389b (DYS389II minus DYS389I). **b.** Comparison of TMRCAs based on Y-SNP and 15 Y-STRs (DYS385a and DYS385b were also not used) using four different Y-STR mutation rates in 78 East Asian samples of Li Jin lab, with the dates estimated based on sequence data. For more details, see supplementary table.2

We next took two ways to see whether the time estimation using genealogical Y-STR mutation rates really corresponds best with sequence-based dating for Neolithic coalescent sublineages. In our previous work, we found three strong star-like Neolithic lineage expansions (Oα, Oβ, and Oγ) at about 5.4-6.8 kya through sequencing 78 East Asian Y chromosomes at 3.9 Mbp of NRY^12^. We used 15 STRs of the 78 individuals to do lineages dating. One evolutionary rate and three genealogical rates are used in Batwing (EMR and rEMR are the same for the 15 STRs we used). The results are very similar with the above analysis using 1000 genome dataset. The sequence-based TMRCAs for Oα, Oβ, and Oγ are almost the same with those estimated using genealogical STR rates, but 3-4 times younger than the times calculated with evolutionary rate. We then validated this result by estimating the TMRCA of 366 individuals belonging to haplogroup Oγ-F11 using 10 STRs (Wang et al., 2013b) in Batwing. This approach is to eliminate the possible bias in time estimation due to small sample size. The TMRCA of Oγ using genealogical rates is around 10 kya, however, TMRCA with evolutionary rate is even more than 30 kya (EMR: median=34.1, mean=46.0, sd=15.6; OMRB: median=11.1, mean=13.9, sd=3.29; OMRS: median=9.30, mean=11.7, sd=2.50; lmMR: median=9.40, mean=12.4, sd=3.87 in kya). The TMRCAs using genealogical rates are more close to that estimated with our previous sequencing data.

We have noticed that TMRCAs for main lineages show large gaps with both the times estimated using evolutionary and genealogical STR mutation rates. This phenomenon reminds us that the evolutionary rate (6.9E-4 per locus per generation) might be too low and the genealogical rates might be too high in for time estimation of main lineages. As the genealogical rates are calculated from multiple pedigrees, each marker has an individual mutation rate, ranging from 4.0E-4 to 1.6E-2 per locus per generation. There might be some Y-STRs lead to more reliable estimates for the above main lineages. We then classified the 21 STRs of 1000 genome samples into two subsets according to their mutation rates: the first ten markers with higher rates are assigned as fast markers, the last ten markers with lower rates are assigned as slow markers (DYS389b was exclude in the analysis). We redid the time estimation in Batwing using fast and slow markers, respectively. The TMRCAs using fast markers (fig.3a) show a very similar pattern with those using combined markers (fig.2a), but the times using evolutionary rates are higher than previous estimates. However, the TMRCAs using slow markers fit well with sequence-based estimates (fig.3b) and are also highly correlated (OMRB: Pearson’s r=0.898, Spearman’s rho=0.956, p= 3.365E-7; OMRS: Pearson’s r=0.870, Spearman’s rho=0.940, p= 1.878E-6) in main lineages.

**Fig. 3.**
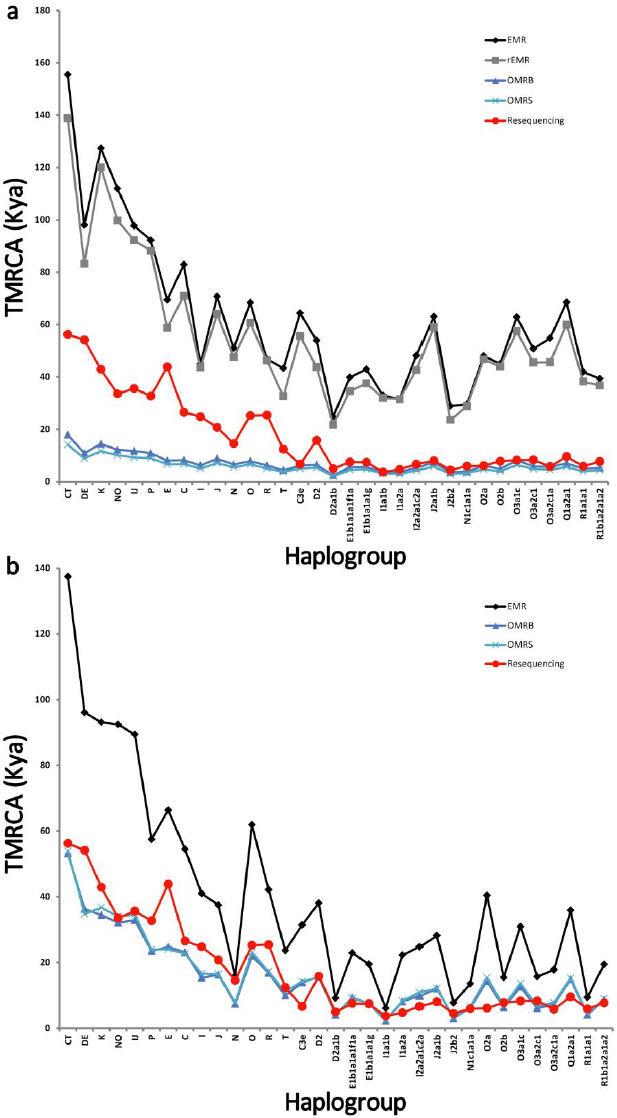
**a.** Comparison of TMRCAs based on Y-SNP and 10 fast mutated Y-STRs using four different Y-STR mutation rates in 377 samples of 1000 genome project, with the dates estimated based on sequence data. **b.** Comparison of TMRCAs based on Y-SNP and 10 slow mutated Y-STRs using three different Y-STR mutation rates in 377 samples of 1000 genome project, with the dates estimated based on sequence data. For more details, see supplementary table.2

## Discussion

In this study, we have compared the Y chromosomal lineage dating between sequence data and commonly used Y-SNP plus Y-STR data in Batwing. The TMRCAs using evolutionary Y-STR mutation rates correspond best with sequence-based dating when the lineages include the most ancient haplogroup A individuals. However, the TMRCAs using slow mutated STR markers with genealogical rates fit well with sequence-based estimates in main lineages, such as haplogroup CT, DE, K, NO, IJ, P, E, C, I, J, N, O, and R. Genealogical rates give times that are more similar to sequence-based dating for Neolithic coalescent sublineages, such as R1b1a2a1a2, E1b1a1a1f1a, E1b1a1a1g, Oα, Oβ, and Oγ.

The conclusion drawn from our study is not an omnipotent rule in Y chromosomal lineage dating. First, all the analysis are calculated in Batwing using stepwise mutation model (SMM) for all the STRs. However, Different time estimation methods use different algorithms and assumptions, thus alternative methods probably fit more or less well with sequence data in time estimations. In addition, the best-fit mutation model might vary for different STRs. Second, some specific lineages might have their own unique best-fit STR mutation rates for time estimation. For instance, TMRCAs for many main lineages show large gaps with both the times estimated using evolutionary and genealogical STR mutation rates. However, the TMRCA for haplogroup E is about 43.8 kya, which is more consistent with the time estimated using evolutionary rates (fig.2a).

## Acknowledgements

This work was supported by the National Excellent Youth Science Foundation of China (31222030), National Natural Science Foundation of China (31071098, 91131002), Shanghai Rising-Star Program (12QA1400300), Shanghai Commission of Education Research Innovation Key Project (11zz04), and Shanghai Professional Development Funding (2010001).

